# Short-time fractal analysis of biological autoluminescence

**DOI:** 10.1101/578286

**Authors:** Martin Dlask, Jaromír Kukal, Michaela Poplová, Pavel Sovka, Michal Cifra

## Abstract

Biological systems manifest continuous weak autoluminescence, which is present even in the absence of external stimuli. Since this autoluminescence arises from internal metabolic and physiological processes, several works suggested that it could carry information in the time series of the detected photon counts. However, there is little experimental work which would show any difference of this signal from random Poisson noise and some works were prone to artifacts due to lacking or improper reference signals. Here we apply rigorous statistical methods and advanced reference signals to test the hypothesis whether time series of autoluminescence from germinating mung beans display any intrinsic correlations. Utilizing the fractional Brownian bridge that employs short samples of time-series in the method kernel, we suggest that the detected autoluminescence signal from mung beans is not totally random, but it seems to involve a process with a negative memory. Our results contribute to the development of the rigorous methodology of signal analysis of photonic biosignals.

## Introduction

Practically all organisms perpetually generate weak light (300–700 nm wavelength range), too weak to be visible to naked human eye, in the course of their internal metabolic processes [1]. This light phenomenon differs from a rather bright bioluminescence which is dependent a specific enzymatic complexes which are present only in very specific species such as fireflies and selected jellyfish. What differentiates the general biological autoluminescence from ordinary bioluminescence is, apart the weaker intensity, its ubiquity across biological species ranging from microorganisms [2–5] through tissue cultures [6–8], plants [9–13] up to animals [14] including human [15–17]. There are also various synonyma used in the literature describing this light phenomenon such as ultra-weak photon emission [18], ultra-weak bioluminescence [19], endogenous biological chemiluminescence [20], biophotons [21–23], etc.

Widely accepted underlying mechanism which generates biological autoluminescence (BAL) is related to a chemical generation of electron-excited states of biomolecules in the course of oxidative metabolism and oxidative stress [18, 24]. While intensity and optical spectrum properties of BAL as a factor of various influences have been widely investigated [3, 25–29], there is a limited knowledge and consensus about statistical properties of BAL.

The object of our current study is the BAL time series from the seeds of mung beans that were measured using a sensitive photomultiplier setup. We decided to test the hypothesis if the BAL signals of mung beans contain any intrinsic correlations. To that end, we recorded and analyzed the time series of the BAL from mung beans. One of the ways to assess correlations in the signal employs chaos- and fractal-based approaches [30]. We focus here on the analysis of the fractal character of time-series using fractional processes.

Fractional Brownian motion (fBm) and fractional Gaussian noise (fGn), introduced by Mandelbrot [31], have been intensively investigated over the last few decades. They are both dependent on Hurst [32, 33] exponent *H* ϵ (0; 1) that influences their autocovariance structure. The fBm or fGn assumption of finite sample is advantageously used in many fields of research of time series analysis – in network traffic modelling [34, 35], financial time series [36, 37], or in biomedicine especially for detection of Alzheimer’s disease [38] and cardiology [39].

When analyzing real-world data, the measured sample is usually discrete and short. The traditional methods are generally not suitable for short time series analysis. That is the reason why we need to use a precise method that can estimate the Hurst exponent without bias and can determine the confidential intervals of the estimate. The fractional character of data can be measured via fractional Brownian bridge model, which is a derived discrete process from traditional continuous fBm. A lot of time series are short due to their nature or cut by purpose or experimental limitations. Reconsidering some fBm properties that are taken in long time series analysis as granted and customizing them into a short-time, the discrete model allows estimating Hurst exponent of the discrete measured signal. This approach is advantageously used in a recently developed method of fractional Brownian bridge [40].

The article at first analyzes current open questions of statistical properties of biological autoluminescence. In the next section, we then describe the theory of fBm and the method of Hurst exponent estimation as well ass other employed methods, whereas the last section contains the results of the analysis of experimental signals compared to computer generated reference signals.

## Statistical properties of biological autoluminescence (BAL)

### Rationale for the need of understanding of BAL statistical properties

Multiple authors proposed that statistical properties of BAL time series might contain an information related to the state of biological system [41–43]. If the existence of such nontrivial statistical properties was rigorously confirmed, it would make a substantial impact on three major areas of this research field.

At first, the discovery of nontrivial statistical properties of BAL would have an impact on the understanding of the BAL generating mechanisms [18, 21]. So far, well accepted generating mechanism of BAL [18, 24] implicitly considers BAL a weak endogenous biological chemiluminescence formed as a by-product of oxidative metabolism and oxidative stress. General chemiluminescence is typically considered to be random arising from individual uncorrelated photon emitter molecules [44]. If any correlations in the signal were observed, one would start to ask questions what physical, chemical, and biological processes generate such correlations, hence casting the light on BAL generating mechanisms.

At second, nontrivial statistical properties might revive an interest into long-standing intriguing, yet unresolved question: does BAL enable optical communication between cells and organisms [45–48]. Underlying hypotheses for such biocommunication role of BAL usually expect that BAL carries information which can be processed by a receiver [46]. Such information could be encoded in the intensity and optical spectrum of BAL [49] or in statistical properties of BAL, if they are any different from random light, as claimed by some authors [45].

At third, statistical properties would represent a completely novel fingerprint for application of BAL in biosensing in biotechnology, agriculture, food industry, and medicine beyond the intensity and optical spectra, hence greatly enhancing application potential of BAL analysis.

### Approaches for analysis of BAL statistical properties

#### Quantum optics approach

Historically, the first common approach to analyze the statistical properties of the photon signals is based on quantum optics theorems and employs photocount statistics of detected photonic signal [50]. Using this approach, several authors suggested that BAL manifests quantum optical coherent properties [21] or even interpreted the observed photocount statistics in terms of quantum optical squeezed states [22, 51]. We have recently criticized the interpretation of experimental evidence claiming quantum optical and quantum coherence properties of BAL [23].

#### Fractal- and chaos-based signal analysis approach

We believe that it is more realistic to consider that BAL could manifest complex statistical or correlated behavior due to the nature of underlying chemical reactions [52] instead of a hypothetical biological coherent quantum field as proposed in the earlier approach. For the analysis of such complex statistical or correlated behavior, fractal or chaos-based methods seem to be appropriate. Therefore, more recent efforts in the analysis of BAL statistical properties were focused on the various measures quantifying the complexity and correlations in the time series such as Hurst exponent [53] and multifractal spectra [54].

Several works found correlations or deviations from purely random process with a trivial properties in the BAL signal [42, 43, 54]. However, in all those cases, either signals of different signal-to-noise ratio [42, 43, 54] or surrogate (randomly reshuffled time series) [43] were used as reference signals. Comparing BAL signals having different signal-to-noise (signal = net mean intensity of BAL, noise = mean value of detector noise) ratio may lead to results indicating different statistical properties due to a trivial fact: statistical properties of experimentally detected BAL signal are formed by a convolution of detector noise properties with a pure BAL properties. We demonstrated this issue on the example of Fano factor analysis in [13, Fig.4].

Using surrogate signals might also lead to misleading interpretation in case the signal contains a certain trivial linear trend before random reshuffling – such reshuffling would eradicate any trend. We showed recently that detrending of the BAL signal is not sufficient to remove artifacts since the trend is present not only in the local mean but also in the local variance of the signal [53, Fig.1b, Fig.4b]. We suggest that the most reliable testing of the hypothesis of non-trivial correlation properties so far can be obtained using reference signals with well-defined properties. To that end, in our recent works, we used computer-generated Poisson signal time series superposed on the experimentally detected detector dark count times series as the control signals with signal-to-noise ratio same as the experimentally detected BAL signals [53]. Such a method for reference signal generation was also recently used in entropy analysis of BAL from model plant *Arabidopsis Thaliana* and helped to correctly interpret findings of different entropy values at different stages of seed germination [20, Fig.6].

For the first time, we combine here the advanced approach of computer generated reference signals [53] and a novel method based on fractional Brownian motion analysis [55] to test if BAL signals from mung beans manifest any correlations.

## Materials and Methods

### Experimental

#### Preparation of Samples

Mung bean seeds (*Vigna radiata*, BIO Mung, CZ-BIO-001) were used as a biological material. Mung seeds were surface-sterilized with 70% ethanol for 1 min. Then, the ethanol was removed and 50% disinfecting agent (SAVO, CZ) was added. After 10 min, the seeds were washed with distilled water 6 times and soaked for 6 h (shaken every half an hour). After the preparation, the green covers of the seeds were removed. Then, they were germinated in dark condition on large Petri dishes with ultra-pure water.

#### Luminescence measurement system

We used a measurement system based on cooled (−30 °C) low-noise photomultiplier tube (PMT) R2256-02 (all components of the system from Hamamatsu Photonics Deutschland, DE, unless noted otherwise), see Fig. 2. Cooling unit C10372 (Hamamatsu Photonics Deutschland, DE) consisted of a control panel and a housing in which the PMT is placed. External water cooling is used for lower cooling temperature. High voltage power supply PS350 (Stanford Research Systems, USA) was used for powering the PMT. C9744 unit, consisting of a preamplifier, discriminator and shaping circuit, transforms photocount pulses coming from the PMT into 5V TTL pulses detected by C8855 unit connected to PC. Discriminator level was set to −500 mV and high voltage PMT supply to –1550 V based on the experimental SNR (signal-to noise-ratio) optimization procedure performed in [56].

The PMT had a dark count of ca. 17.2 s^*−*1^ and photocathode diameter 46 mm); see its quantum efficiency in Fig. 2. PMT was mounted from the top outer side of the black light-tight chamber (standard black box, Institute of Photonics and Electronics of the CAS, CZ). The distance between the PMT housing input window and the inner side of the bottom of the Petri dish was 3 cm.

#### Measurement protocol

The second day after the preparation day, 12 similar mung beans were chosen for the study and distributed into a Petri dish (5 cm in diameter), see Fig. 2.

### Short Sequence Analysis

#### fBm Hypothesis

Fractional Brownian motion (fBm) [31] is a continuous Gaussian process *B_H_* (*t*) defined for continuous variable *t* ∈ [0; +∞), *H* ∈(0; 1) and *σ >* 0. The process starts at zero and has zero expected value for all positive times *t*. The autocovariance structure of fBm obeys for all *t, s >* 0

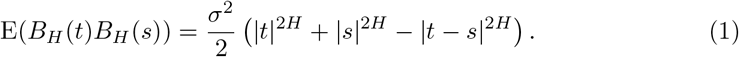

Parameter *H* is called Hurst exponent, for *H* = 1*/*2, the fBm becomes Wiener process, which is standard Brownian motion. There are several cases of time series behaviour:

- *H →* 1^*−*^ as strongly dependent and predictable,
- *H ∈* (1*/*2; 1) as positive long memory process,
- *H* = 1*/*2 as Wiener-like process,
- *H ∈* (0; 1*/*2) as negative long memory process,
- *H →* 0^+^ as strongly dependent, but hardly predictable.

Discrete fractional Brownian motion of length *N ∈* N is any discrete process defined for discrete variable *k* = 0*,…, N −* 1 with zero mean and autocovariance function defined for *k, l* = 0*,…, N −* 1 and *l < N − k* as

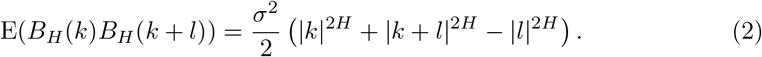

Taking a sample of fractional Brownian motion, it is possible to investigate short samples of time series with fractional character. Finite sample *B*_*H*_ (*k*) of size *N* + 1 for *k* = 0*,…, N* of standardized fBm can be used for the construction of fractional Brownian bridge [55] in the following way

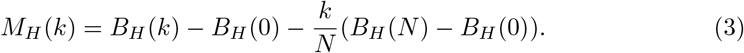

In the fractal analysis of time series, the fractional processes are often converted to fractional noises utilizing signal difference to simplify their covariance structure together with its spectral properties keeping the desired dependence on Hurst exponent. The differenced fractional Brownian bridge (dfBB) [55] is defined as

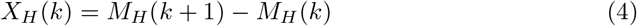

for *k* = 0*,…, N −* 1.

#### Theory of dfBB

The dfBB is a discrete process and it is proven that the process has zero expected value and its variance is independent on the time lag and equals

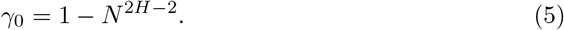

The autocovariance of dfBB can be expressed as

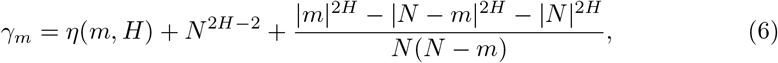

for *m* = 0, 1*,…N −* 1 where

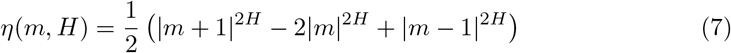

The corresponding autocorrelation function is again independent on the time lag and can be expressed as

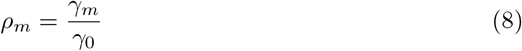

for *m* = 0*,…, N − 1*. The autocorrelation function of dfBB for selected *H* is depicted in Fig. 1.

The estimation of Hurst exponent will be based on the correlation function (8). This correlation function is valid only for discrete processes that originated as sampling continuous fBm. In our work, we assume that the investigated signals have the fBm property with unknown Hurst exponent. The advantage of using dfBB is the de-trending of the input signal which is important in the real experiment outcome analysis.

**Fig 1.**
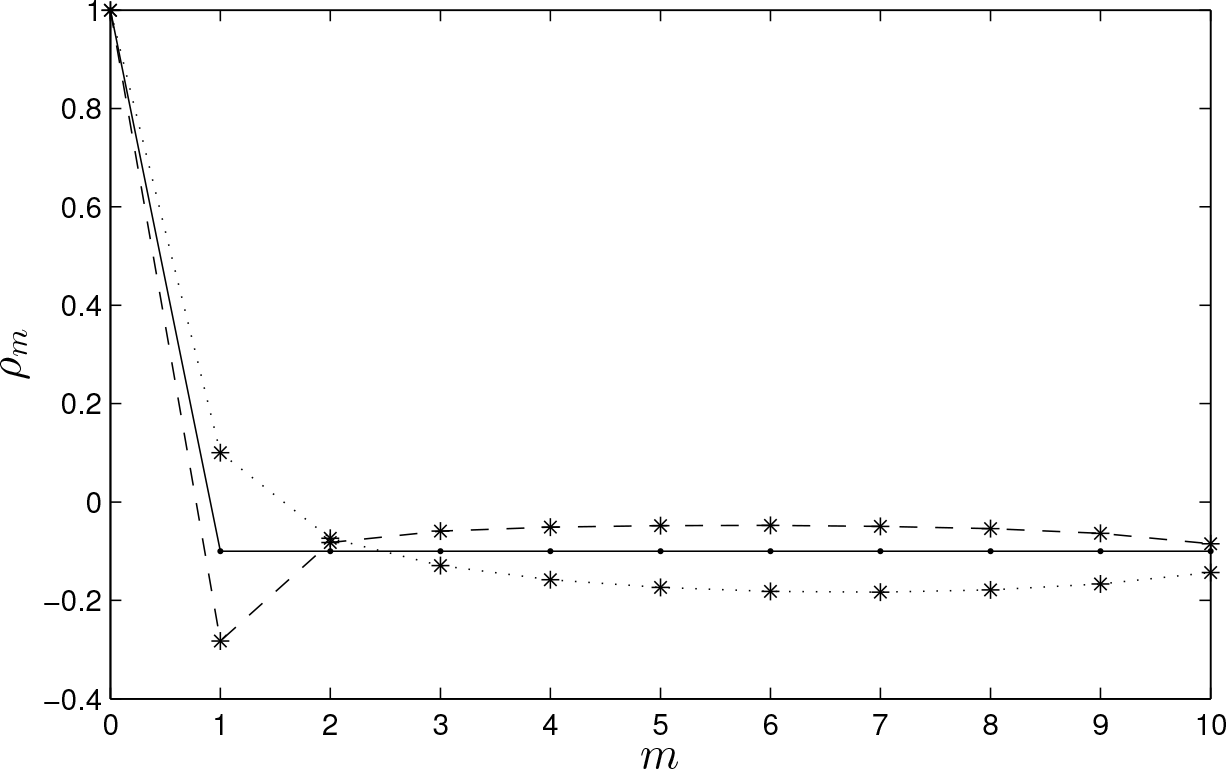
Autocorrelation function of dfBB for *H* = 0.7 (dotted), *H* = 0.3 (dashed) and *H* = 0.5 (solid line).

**Fig 2.**
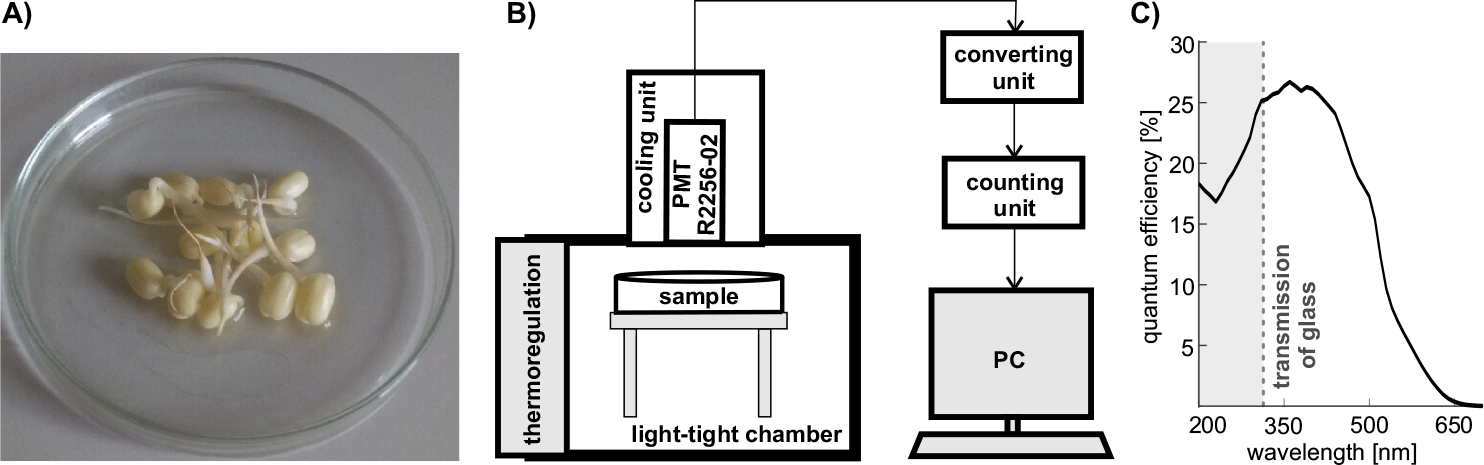
A: Sample of germinating mung beans. B: Scheme of the luminescence measurement setup. C: Quantum efficiency of the photomultiplier used for the detection of biological autoluminescence

### Hurst Exponent Estimation

The estimation of Hurst exponent is based on the fitting of the autocorrelation function. For an input discrete signal that has the fBm properties, the dfBB according to formulas (3), (4) is created. If the original signal has length *N* + 1, the respective dfBB has length *N* having elements *x*_0_*, x*_1_*,…, x*_*N*−1_. The estimation of *n*-th autocovariance coefficient 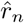 can be expressed for *n* = 0*,…, N −* 1 as

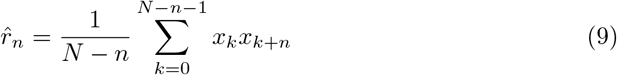

in the case of unbiased estimation. Alternative biased estimate is based on formula

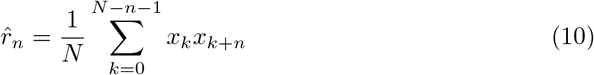

and the estimation of autocorrelation coefficient 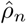 as

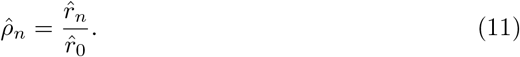

The results using (9) and (10) were proven to be comparable, therefore we used the equation (9) for the following calculations. Denote the theoretical value of autocorrelation from equation (8) as *ρ*_*n*_ = *ρ*_*n*_(*H*) and the experimentally calculated autocorrelation as 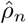. Than we obtain the estimation of parameter *H* by means of solving the minimization problem

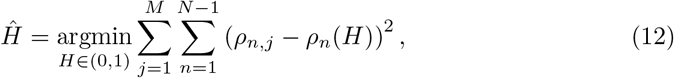

where *M* is the number of signal segments. The point estimate of 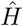 was obtained by the maximum likelihood method [57] together with its standard deviation *ŝ* as recommended in [55].

#### Reference signal generation

Recently we demonstrated that a suitable reference signal is crucial to understand and interpret the findings from various BAL signal analysis [20, 53]. Detector noise itself is not a suitable reference signal since it contains intrinsic technogenic correlations itself [53] and using signals of other samples with different signal-to-detector noise ratio can also lead to misleading results as we explained in section. Hence for this work we follow our method [53], and generated the reference signal as a sum of measured detector noise and computer-generated Poisson signal (using Matlab®2017 *poissrnd* command) with given *λ* in every experimental point where *λ* = E*λ*_MUNG_ - E*λ*_NOISE_. The respective values of *λ* in case of 200 µs signal as well as 500 µs signal are calculated in Tab. 2. Hence, experimentally detected BAL signals from mung beans and reference signals have practically the same mean value and same signal-to-noise ratio.

## Results

### Measurement

The investigated sample of germinating mung beans is displayed in the Fig. 2. An overview of all signals collected and employed in this paper is in Tab. 1.

There were two bin size settings used to collect the signals: *T*_s_ = 200 and 500 µs. For each sampling period, we have corresponding mung bean signals, detector noise signals and computer generated reference signals.

Both mung beans signal and PMT detector noise signal are assumed to be stationary with their mean values with Poisson distribution. Therefore, they can be represented by their mean values E*λ*_MUNG_ and E*λ*_NOISE_ that are estimated from the measured data.

As previously mentioned, the aim of study is to compare mung beans signal with the reference signal and find statistical difference between them using their autocorrelation. With each of these two signals independently, we performed basic data processing. This procedure describes the normalization of the data, which is the essential property of fBm processes. At first the input time series *y*_*k*_ for *k* = 0, 1*,…, Q −* 1 was cumulatively summed for a window size 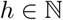 and Anscombe transformation [58] was performed. The resulting signal *z*_*k*_ can be expressed based on the output from measuring device *y*_*k*_ as

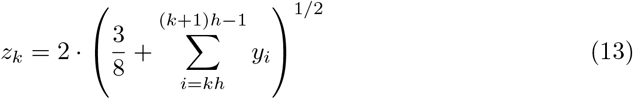

for *k* = 0,…, *M* 1. This transformations assures stationarity by terms of variance and guarantees Gaussian distribution of the resulting signal.

### Likelihood ratio test

Having signal from the mung beans photon emission as well as the reference signal, we will use likelihood ratio test [59] to decide, whether the Hurst exponent of both samples is significantly different. We denote *H*_D_ as the Hurst exponent estimate of the PMT detector noise or reference signal and *H*_B_ as the Hurst exponent estimate of mung emission using the formula (12). The overall error (sum of the squares of residuals) is defined as

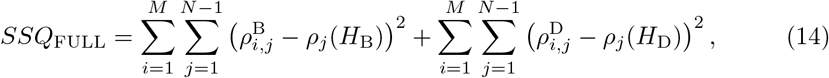

where *ρ*^D^, *ρ*^B^ are the autocorrelation coefficient of the noise and photon emission, respectively. The case of *j* = 0 is excluded due to 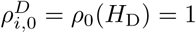 for all *i* = 1*,.., M*. Using sub-model satisfying *H*_B_ = *H*_D_ we get

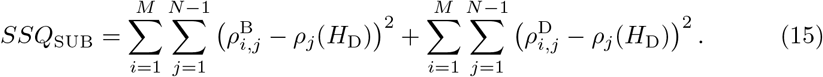

Using likelihood ratio (LR) test of significant difference between the sub-model and the full model, we calculate

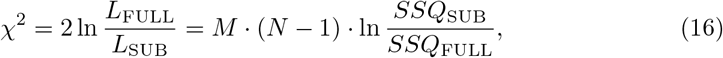

where *L*_FULL_ and *L*_SUB_ are corresponding likelihoods.

When the hypothesis H_0_ : *H*_D_ = *H*_B_ holds, i.e. the full model has the same validity as the submodel, the criterion has 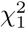 distribution due to single parameter constrain.

### Hurst Exponent Estimates

There is no prior knowledge of optimal model length, segment length, and Hurst exponent. Therefore, we will apply the maximum likelihood method of Hurst exponent estimation for the various model and segment lengths, and then we will individually test the differences in the Hurst exponent. However, there is a finite number of reasonable pairs (model length *N*, segment length *h*) which will cause the phenomenon of the multiple hypothesis testing. After the False Discovery Rate (FDR) correction, we will localize the model and segment lengths which causes significant differences in the Hurst exponent. These pairs (*h, N*) will be declared as significantly sensitive to the signal differences in the Hurst exponent.

Having signals with two different bin sizes, we will use the signal bin size *T*_*b*_ = 200 µs as training set and the signal with *T*_*b*_ = 500 µs as a verification set. Normalized mung beans and reference signals with bin size *T*_*b*_ = 200 µs and length *Q* = 500 000 were the subject of the initial analysis. The bin compression of size *h* was applied to the signals, therefore the number of bins was [*S/h*]. After the bin compression, the signal is divided into segments of length *N*. Due to the memory of fBm process, we will use only the odd segments for the calculation of autocorrelation function and the even segments are excluded. The new signal has length *⌈⌊⌊Q/h⌋/N⌋/*2⌉. Using equation (12) and maximum likelihood method, we obtain the corresponding *H*_D_ and *H*_B_ estimates for the Hurst exponent of referential signal and mung beans, respectively. Based on these estimates, we can derive the *p*-values of LR test using (16) statistics.

In our case, we performed altogether 11 × 11 = 121 tests for *h* = 1500, 1550,…2000 and *N* = 20, 21,…30. Due to multiple testing and obeying the Hochberg-Benjamini principle, we diminish the significance level from 0.05 to *α*_FDR_ = 0.000050. The *p*-values as decadic logarithms are shown in Tab. 3.

In these settings, there were two cases where the Hurst exponent was significantly different. The results from these two cases are displayed in Tab. 4. The lowest *p*-value was obtained in the case of (*h, N*) = (1750, 24), which represents the segmentation into bins with duration 1750 *×* 200 µs = 350 000 µs = 0.35 sec.

As the verification set, the signal with *T*_s_ = 500 µs was taken into account, following the same procedure as the previous one. The bin compression *h* was accordingly diminished to 2*/*5 of its previous value to guarantee the same segment length. Initially, we have three types of signal:

- (B) - mung beans signal,
- (D) - noise signal of PMT detector,
- (R) - reference signal as a sum of measured detector noise (D) and computer-generated Poisson noise.

We perform the verification for the combination of signals (B) and (R) similarly as in the previous case and additionally for the combination of (B) and (D). The first set of signals ((B) and (R)) will be used to test if the photon emission is not random and has a negative memory, while the results from the second set ((B) and (D)) of signals will be used to test if there is a significant difference between the cases, when the PMT detects BAL signals from mung beans compared to PMT noise. We use the significant cases from Tab. 4 to estimate their Hurst exponent and the results on verification set is displayed in Tab. 5. The variables *s*_1_, *s*_2_ denote the pair of signals, whereas the *H*_X_ denotes the estimation of Hurst exponent of the signal *s*_2_.

We performed 4 tests and according to Hochberg-Benjamini false discovery rate, we diminish the *α*_FDR_ = 0.0169. Therefore, all four cases are considered significant and we reject the hypothesis that the Hurst exponent of mung beans would be the same as *H*_X_.

## Discussion

Results from rigorous statistical analysis and testing in tables 3, 4, and 5 suggest that the mung beans signal has a negative memory (negative correlations, antipersistent behavior [60]) and its Hurst exponent is lower than the referential signal. How could such behavior originate in biological systems ? It was proposed that the restriction of Brownian motion due to structuring of nano- to microscale intracellular environment leads to anomalous sub-diffusion [61] characterized by Hurst exponent *<* 0.5 [60]. This is understandable since a cytoplasm environment displays fractal spatial structuring [62]. Since biochemical reactions (encounters of reactants) leading to BAL are taking place within the cell cytoplasm, organelles and lipid membranes [24] where anomalous sub-diffusion was observed [61, 63], it is not a great logical leap to speculate that BAL from mung bean samples could also display sub-diffusive features. Actually, it is already acknowledged that chemical reactions spatially constrained on the microscopic level may lead to fractal reaction kinetics [64–66] also in case of intracellular biochemical kinetics [67]. The 0.35 s as the time scale where we found statistically significant differences of mung bean signal Hurst exponent from that of the reference signal (Tab.3) could correspond to a rate of underlying rate-limiting step of chemical reactions or processes which gives rise to BAL. However, one has to be careful in the interpretation since there are many pitfalls in an accurate estimation of the Hurst exponent value from experiments [68, 69]. Although unlikely, given the nature of our experiments we can not fully exclude that the correlations we observe in mung signals are introduced by the photodetector (PMT) due to the nature of photocounting process [70, 71]. Introduction of anti-/correlations could be at the physical level of the PMT tube (after-pulsing, a temporary drop of the voltage at dynodes after ejecting electrons,…) or the follow-up circuitry (amplifiers). Anti-correlations of the detected counts depending on the count rate have been actually observed due to a PMT construction [71, Fig.9]. However, marked anti-correlations were present only for very high count rates (*>* kHz) and very low quantum efficiency which is not the case in our experiments. We also believe that the dead-time of a PMT [72] is not affecting the value of correlations we observe since the PMT dead-time is on the time scale of few hundreds of nanoseconds - several orders of magnitude smaller than the time scale of correlations we observed (0.35 s) and three orders of magnitude smaller than our bin size (200 and 500 µs).

**Table 1.** Number and type of the signals collected.

| bin size | | 200 $\mu\text{s}$ | 500 $\mu\text{s}$ |
| --- | --- | --- | --- |
| signal type | mung beans (B) | N=5 | N=5 |
|  | detector noise (D) | N=5 | N=5 |
|  | reference signals (R) | N=5 | N=5 |
| number of bins in each measurement | | $N_b=100\ 000$ | $N_b=100\ 000$ |
| length of each measurement [s] |  | 20 | 50 |
| total number of bins $Q=N \times N_b$ | | 500 000 | 500 000 |
| total length of all measurements per signal type [s] |  | 100 | 500 |

**Table 2.** Mean values of mung beans signal and noise.

| $T_b$ | 200 $\mu\text{s}$ | 500 $\mu\text{s}$ |
| --- | --- | --- |
| $E\lambda_{\text{MUNG}}$ | 0.0115 | 0.0288 |
| $E\lambda_{\text{NOISE}}$ | 0.0036 | 0.0088 |
| $\lambda$ | 0.0079 | 0.0200 |

**Table 3.** Difference between the estimated Hurst exponent of mung beans and reference signal as (*−*log_10_*p*)-values.

| $h \setminus N$ | 20 | 21 | 22 | 23 | 24 | 25 | 26 | 27 | 28 | 29 | 30 |
| --- | --- | --- | --- | --- | --- | --- | --- | --- | --- | --- | --- |
| 1500 | 1.188 | 2.934 | 1.835 | 1.888 | 3.175 | 1.645 | 2.284 | 0.863 | 0.506 | 0.192 | 0.762 |
| 1550 | 1.172 | 1.617 | 0.394 | 0.887 | 1.420 | 1.470 | 0.912 | 2.113 | 1.651 | 0.026 | 0.691 |
| 1600 | 1.978 | 1.576 | 0.646 | 0.523 | 1.217 | 0.394 | 1.597 | 0.786 | 1.487 | 0.880 | 1.859 |
| 1650 | 0.990 | 1.616 | 1.127 | 2.024 | 1.209 | 0.651 | 1.635 | 0.909 | 1.906 | 3.573 | 2.927 |
| 1700 | 0.772 | 0.621 | 1.288 | 1.196 | 1.239 | 0.488 | 0.407 | 1.175 | 2.658 | 0.463 | 0.776 |
| 1750 | 1.475 | 2.325 | 1.269 | 3.131 | **4.638** | 1.535 | 2.370 | 1.017 | 0.726 | 0.412 | 1.945 |
| 1800 | 0.465 | 1.455 | 1.394 | 1.098 | 1.313 | 0.180 | 2.661 | 2.064 | 2.449 | 1.917 | 2.001 |
| 1850 | 2.377 | 2.010 | 1.308 | 0.567 | 1.533 | 2.382 | 3.184 | **4.301** | 3.328 | 2.418 | 1.968 |
| 1900 | 2.599 | 0.879 | 0.850 | 0.629 | 1.053 | 1.264 | 0.950 | 0.943 | 1.397 | 2.093 | 0.142 |
| 1950 | 2.574 | 0.095 | 0.706 | 1.900 | 2.843 | 2.874 | 3.261 | 2.514 | 3.462 | 2.501 | 2.405 |
| 2000 | 2.212 | 1.611 | 1.315 | 0.935 | 1.040 | 1.232 | 0.922 | 0.282 | 0.366 | 1.159 | 0.963 |

**Table 4.** Estimated Hurst exponent values for mung beans (B) signal and reference signal (R).

| $h$ | $N$ | $H_B$ | $H_R$ | $p$-val | $-\log_{10} p$-val |
| --- | --- | --- | --- | --- | --- |
| 1750 | 24 | 0.4142 | 0.5299 | $2.3135 \times 10^{-5}$ | 4.638 |
| 1850 | 27 | 0.3569 | 0.4291 | $4.9977 \times 10^{-5}$ | 4.301 |

**Table 5.** Estimated Hurst exponent values from verification dataset. *h*=700 for 500 µs signals corresponds to *h* =1750 for 200 µs signals.

| $s_1$ | $s_2$ | $h$ | $N$ | $H_B$ | $H_X$ | $p$-val |
| --- | --- | --- | --- | --- | --- | --- |
| B | R | 700 | 24 | 0.4032 | 0.4415 | 0.0130 |
| B | R | 740 | 27 | 0.3761 | 0.4112 | 0.0042 |
| B | D | 700 | 24 | 0.4032 | 0.4378 | 0.0169 |
| B | D | 740 | 27 | 0.3761 | 0.4480 | 0.0054 |

Throughout the analysis, the lower limit for parameter *h* was chosen as 1500 to assure the normality of the processed data due to the sparsity of the input signal. Higher accumulation than 2000 is not useful since then we would lose the precision of estimate due to the short length of investigated time-series. The minimal length of segment *N* was chosen to assure consistency of the used model, segment lengths of *N >* 30 do not significantly contribute to the higher precision of estimate [40].

## Conclusion

In this work, we focused on statistical properties of biological autoluminescence from germinating mung bean sample. Our emphasis was on the development of a rigorous mathematical and statistical methodology which takes into account proper reference signals, likelihood ratio test and multiple hypothesis testing effects.

We used a highly sensitive photomultiplier-based detection system to record time-series of photon counts of the mung bean sample emission and noise of the detector. Using the normalization of the input signal we were able to employ the fractional models that allowed us to estimate Hurst exponent. Dividing the input signals into the training set and evaluating the differences in the Hurst exponent of both signals, the procedure allowed us to test our initial hypothesis on the verification signal. The resulting Hurst exponent mean value of mung bean sample time series is below the level of 1/2 which confirmed our initial hypothesis, that the biological autoluminescence displays correlations. We also proposed that this value could be related to anomalous sub-diffusive features of biochemical reactions underlying processes within mung beans which give rise to photon emission time series. Further extensive work beyond the scope of this methodical paper needs to be carried out to test the biological ubiquity of anti-/correlations in biological autoluminescence signals and the role of the detector in the observed Hurst exponent values. Especially interesting would be an analysis of BAL statistical properties across samples with rising complexity starting from simple chemical solutions of small biomolecules through isolated cellular structures and cell suspensions up to whole tissues and organisms. Nevertheless, we believe that rigorous methodology we presented here will help to support the future research of BAL statistical properties towards a deeper understanding of BAL mechanisms as well as applications for label-free and non-invasive analysis in medicine and biotechnology using completely new signal fingerprint types.

## Supporting information

S1 data All raw data files used in our analysis. A1502-data.zip

## Author contributions statement

Contribution roles according to CRediT https://dictionary.casrai.org/Contributor_Roles:

**Conceptualization**: Michal Cifra

**Data curation**: Martin Dlask, Michaela Poplová, Michal Cifra

**Formal analysis**: Martin Dlask

**Funding acquisition**: Jaromír Kukal, Michal Cifra

**Investigation**: Martin Dlask, Michaela Poplová

**Methodology**: Jaromír Kukal, Pavel Sovka, Michal Cifra

**Project administration**: Jaromír Kukal, Michal Cifra

**Resources**: Jaromír Kukal, Michal Cifra

**Software**: Martin Dlask, Michaela Poplová

**Supervision**: Jaromír Kukal, Pavel Sovka, Michal Cifra

**Validation**: Michal Cifra

**Visualization**: Martin Dlask, Michaela Poplová

**Writing – original draft**: Martin Dlask, Michal Cifra

**Writing – review & editing**: Jaromír Kukal, Pavel Sovka, Michal Cifra

